# Bruton’s Tyrosine Kinase Supports Gut Mucosal Immunity and Commensal Microbiome Recognition in Autoimmune Arthritis

**DOI:** 10.1101/2021.03.10.434762

**Authors:** Rachel H. Bonami, Christina E. Thurman, Camille S. Westlake, Lindsay E. Nyhoff, Bridgette B. Barron, Peggy L. Kendall

## Abstract

Bruton’s tyrosine kinase (*Btk*) deficiency preferentially eliminates autoreactive B cells while sparing normal humoral responses, but has not been studied in mucosal immunity. Commensal microbes are essential for arthritis in K/BxN mice, used here to examine how BTK-mediated signaling interfaces with the microbiome. *Btk*-deficient K/BxN mice were found to have small Peyer’s Patches with reduced germinal center and IgA^+^ B cells. Although lamina propria IgA^+^ plasma cells were numerically normal, intestinal IgA was low and IgA coating of commensal bacteria was reduced. IgA-seq showed a shift in microbes that are normally IgA-coated into the uncoated fraction in *Btk*-deficient mice. In this altered microbial milieau, the proportion of *Parabacteroides distasonis* was reduced in *Btk*-deficient K/BxN mice. To determine whether *P. distasonis* contributes to arthritis, it was reintroduced into antibiotic-protected K/BxN mice, where it restored disease. This suggests that *P. distasonis’* inability to thrive in *Btk*-deficient mice may be a factor in disease protection. Thus, BTK supports normal intestinal IgA development, with downstream effects on the microbiome that may contribute to autoimmunity.

## Introduction

More IgA is produced daily than all other antibody isotypes combined, highlighting its importance to host health ^1^. IgA-deficient humans are relatively resistant to infection, presumably due to functional compensation by secreted IgM, but they do have a propensity for autoimmunity ^2–4^. The role of B cell signaling in IgA development has not been well-studied. B cells in mucosal lymphoid tissues undergo IgA class-switch and affinity maturation, giving rise to plasma cells that migrate to the small intestinal lamina propria. B2 B cells in Peyer’s patches undergo T-dependent responses in germinal centers (GCs) while B1 cells that traffic from the peritoneal cavity are more likely driven by T-independent responses without requiring GCs ^5–7^. It is thought that commensal recognition by the immune system via IgA occurs primarily through T-independent responses, whereas pathogens more typically elicit T-dependent responses ^6^. This is a reciprocal process as microbes are required for adaptive immune structure formation in the gut; germ-free mice fail to develop normal Peyer’s patch structures or IgA-secreting plasma cells ^8^. Plasma-cell secreted IgA is transcytosed as a dimer into the gut lumen where it can bind to commensal or pathogenic microbes ^1^. IgA coating of microorganisms can block their attachment to host epithelia, thereby helping to maintain barrier function, or alternatively promote their retention in the mucous layer ^9,10^.

B cell functions depend upon signaling through the B cell receptor (BCR). The B cell signaling protein Bruton’s tyrosine kinase (BTK) serves as a “rheostat” in mature B cells, acting to propagate and amplify signals. Elevated BTK phosphorylation has been observed anti-citrullinated protein antibody (ACPA)-positive, but not ACPA-negative rheumatoid arthritis patients relative to healthy controls ^11^, highlighting a link between overzealous BTK signaling and ACPA production. *Btk*-deficiency in mice is known to reduce germinal center formation and T-independent immune responses, as well as autoantibodies, but T-dependent responses can be elicited, and IgG levels are near normal ^12–18^. In humans, BTK is necessary for B cells to mature beyond the bone marrow, making study of this protein in mature peripheral B cells difficult ^19^. Its role in mucosal immunity has not previously been examined, with the exception that serum IgA was initially found to be normal in *Btk*-deficient mice ^14^.

*Btk*-deficiency protects against autoimmune arthritis in K/BxN mice ^12^. This is consistent with a role for *Btk* in autoreactive B lymphocyte development and function, as arthritis is provoked in this model by autoantibodies ^20^. Commensal microorganisms are also central to autoimmune arthritis development, as germ-free or antibiotic-treated K/BxN mice are protected from spontaneous disease development ^21^. Mono-colonization with segmented filamentous bacteria (SFB) drives autoimmune arthritis in this model, confirming commensal bacteria can trigger disease ^21^.

The crosstalk between B lymphocytes and commensal bacteria is incompletely understood. We therefore investigated how *Btk*-deficiency impacts B lymphocyte populations and responses to commensal bacteria in the gut in the setting of autoimmune arthritis. Peyer’s patch germinal centers are dramatically reduced, but lamina propria IgA^+^ plasma cells are preserved in *Btk*-deficient K/BxN mice. These IgA^+^ plasma cells do not show diminished BCR diversity or somatic hypermutation, but IgA-coated bacterial composition is shifted in the small intestine of *Btk*-deficient mice. These data show that impaired B cell receptor signaling shifts the dynamic interplay between host B lymphocytes and commensal bacteria in the gut.

## Results

### IgA-switched and germinal center B cells are reduced in Peyer’s patches of Btk-deficient K/BxN mice

The impact of *Btk* deficiency on B lymphocyte development and function in the gut has not been investigated. Flow cytometry was used to test how loss of *Btk* alters B cell numbers, isotype switch, and differentiation in the Peyer’s patches and lamina propria of the small intestine in the setting of autoimmune arthritis. Numbers of B cells, germinal center B cells, and IgA^+^ B cells were significantly reduced in Peyer’s patches of *Btk*-deficient K/BxN mice compared to *Btk*-sufficient K/BxN littermate controls (Fig. 1). The frequency of Peyer’s patch B cells that differentiated into germinal center B cells or class-switched to IgA was not significantly different between *Btk^+/y^* and *Btk^−/y^* K/BxN mice; the average frequency of total B cells or IgA^+^ B cells that acquired the germinal center phenotype was ~10% and 40%, respectively, regardless of BTK expression (Fig. 1). Taken together, these data suggest *Btk*-deficiency reduces but does not block B cell class switching to IgA or participation in germinal center reactions.

**Figure 1.**
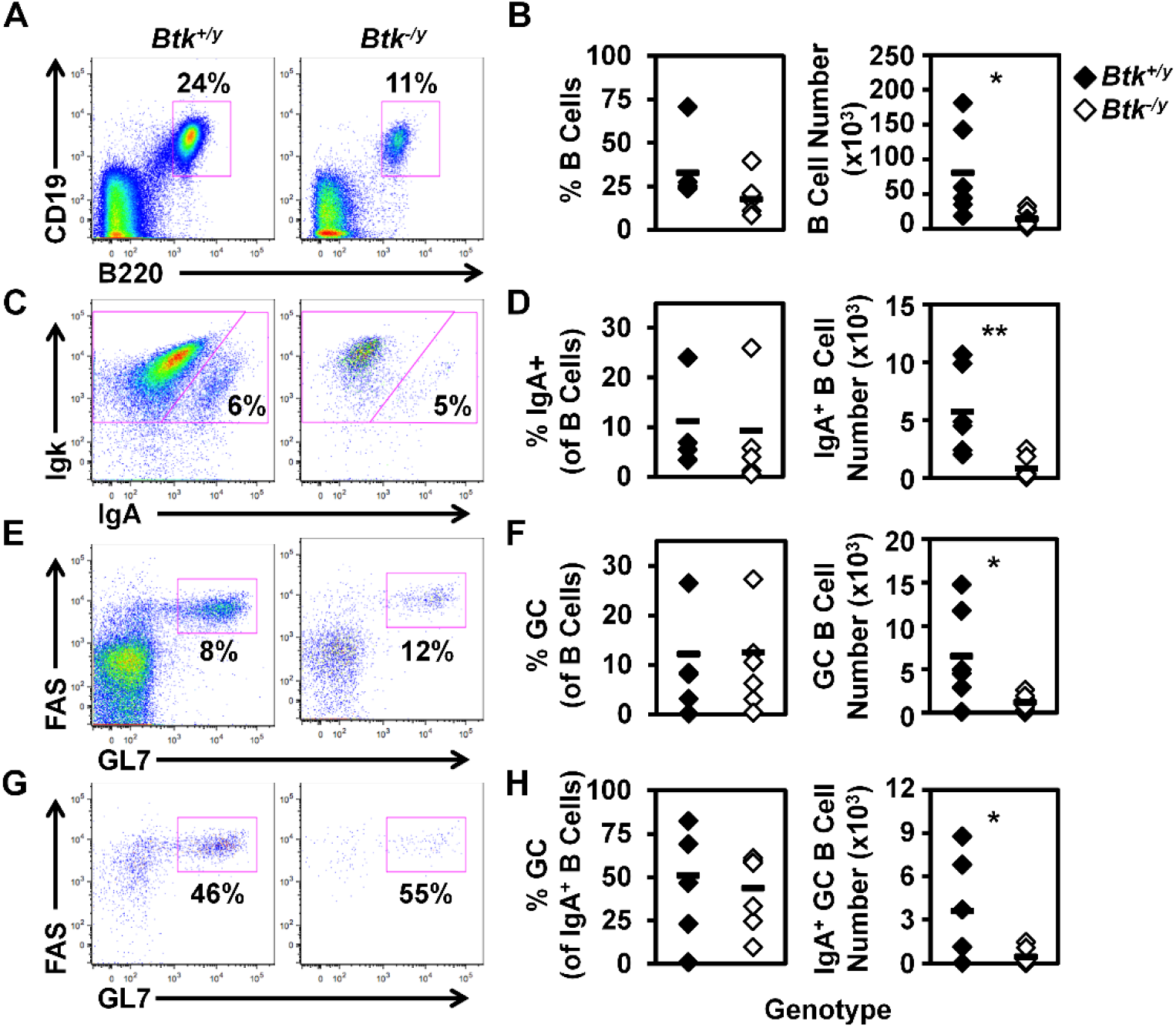
BTK absence leads to reduced numbers of germinal center and IgA class-switched B cells in the Peyer’s patches in K/BxN mice. Peyer’s patches were harvested from K/BxN (black) and *Btk*^−/y^/K/BxN (white) mice as in Methods. Flow cytometry was used to identify the indicated subsets. (A) Representative plots depict B cells in K/BxN (left) and *Btk*^−/y^/K/BxN (right) mice among live lymphocytes. (B) Total B cell frequencies (left) and numbers (right) are shown. (C) Representative plots show IgA+ B cells among B220^+^ CD19^+^ live lymphocytes. (D) IgA^+^ B cell frequencies (left) and numbers (right) are shown. (E) Representative plots show germinal center (GC) B cells among B220^+^ CD19^+^ live lymphocytes. (F) GC B cell frequencies (left) and numbers (right) are indicated. (G) Representative plots show germinal center (GC) B cells among IgA^+^ B220^+^ CD19^+^ live lymphocytes. (H) The frequency (left) and number (right) of GC B cells is shown among IgA^+^ B cells. n ≥ 6 8-9 wks old mice per group, * p < 0.05, ** p < 0.01, *** p < 0.001, two-tailed t test.

### B1a B cells are reduced in Btk-deficient K/BxN mice

IgA-secreting plasma cells in the lamina propria of the small intestine are derived from B2 cells in the Peyer’s patches and B1 cells in the peritoneal cavity ^5,6^. B1a cells are known to be dramatically reduced in *Btk*-deficient mice, while B1b cell numbers are normal ^14^. Consistent with observations in a non-autoimmune strain ^14^, *Btk*-deficient K/BxN mice had a strongly reduced population of B1a B cells in the peritoneal cavity (Fig. 2). Thus, both B1a and B2 B cell plasma cell precursors are reduced in *Btk*-deficient K/BxN mice.

**Figure 2.**
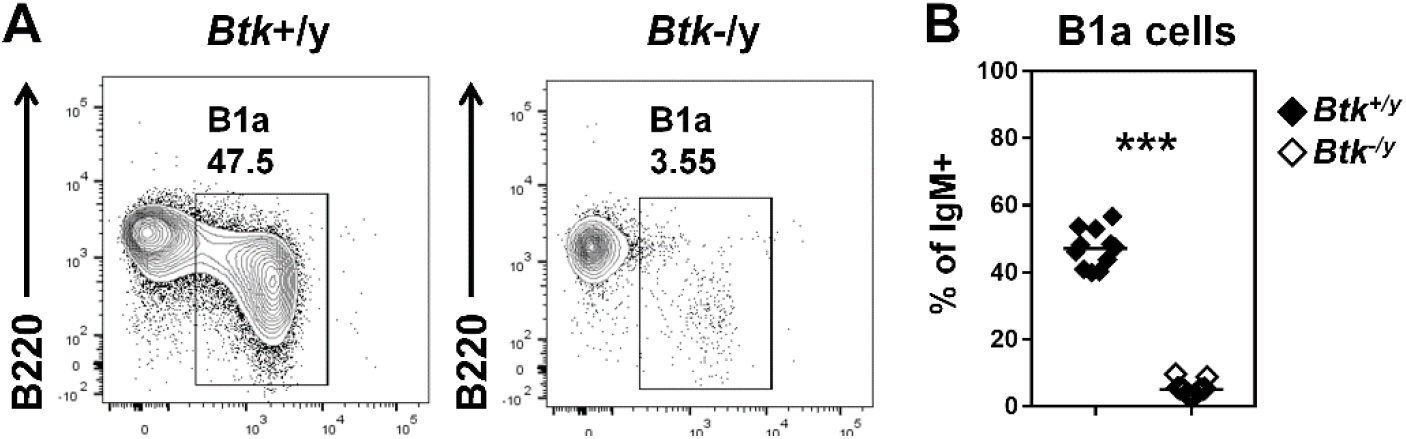
B1a B cells are reduced in *Btk*-deficient K/BxN mice. Flow cytometry identifies B1a B cells in peritoneal lavage of K/BxN (black) and *Btk^−/y^*/K/BxN (white) mice. (A) Representative plots are gated on IgM^+^ live lymphocytes and indicate the frequency of B1a B cells (B220^low^ CD5^+^). (B) n ≥ 11 mice per group are plotted, *** p < 0.001, Mann-Whitney U test.

### IgA^+^ plasma cells persist in K/BxN mice despite loss of BTK

Given the striking reductions in IgA^+^ germinal center B cells in Peyer’s patches and B1a cells which serve as intestinal plasma cell precursors (Fig. 1–2), we sought to determine whether *Btk* loss also reduced IgA^+^ plasma cells in the lamina propria of K/BxN mice. Despite the reduction in Peyer’s patch B cell populations, *Btk*-deficiency did not result in reduced IgA^+^ plasma cell numbers (Fig. 3). The majority of B220^−^ CD19^−^ CD138^+^ plasma cells in the lamina propria were IgA class-switched in both *Btk*-sufficient and deficient K/BxN mice (Fig. 3). Thus, IgA plasma cells are present at normal numbers in the lamina propria despite marked Peyer’s patch germinal center B cell reduction and lack of B1a cells in *Btk*-deficient mice.

**Figure 3.**
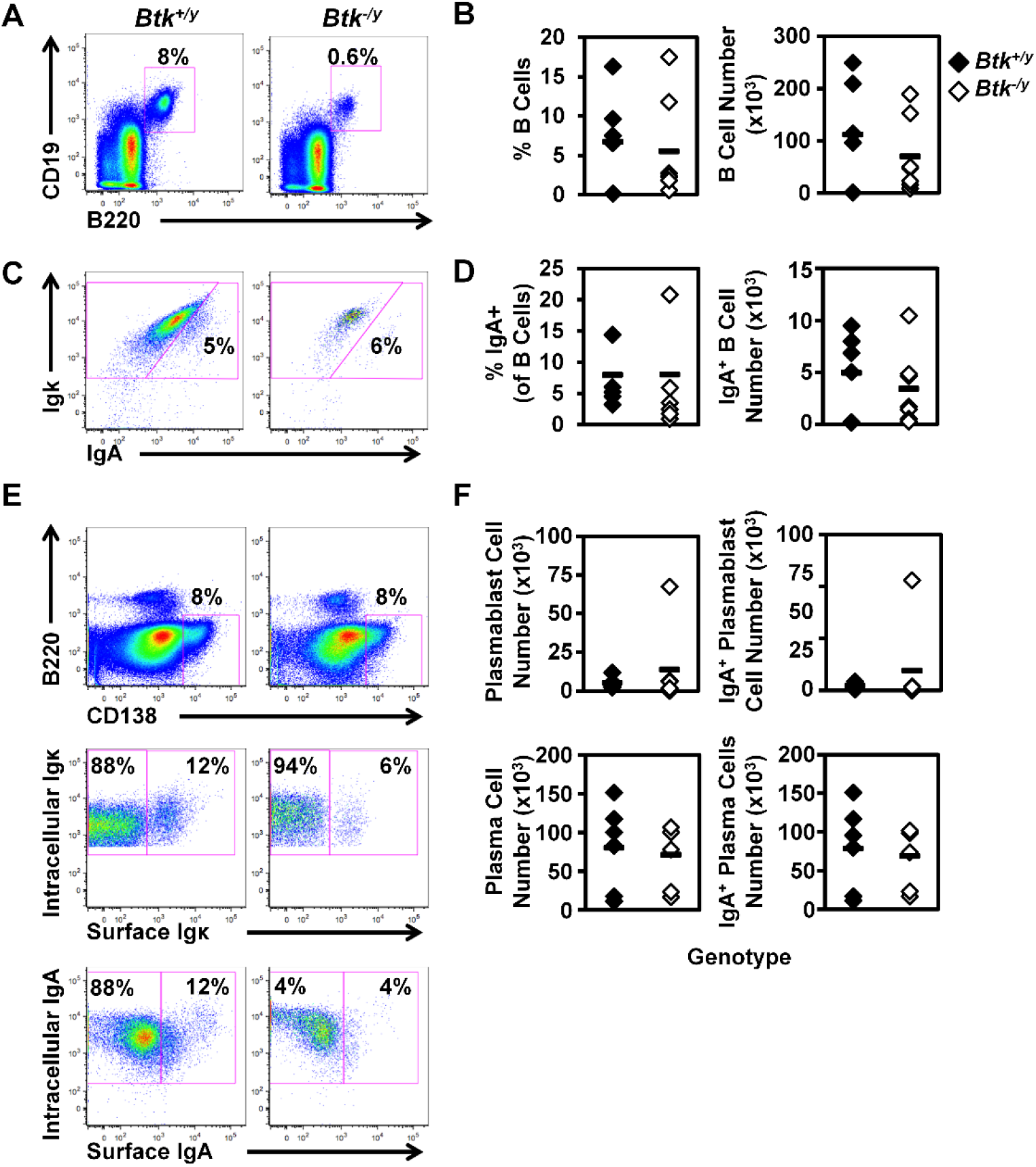
IgA+ plasma cells persist in the lamina propria of *Btk*-deficient K/BxN mice. Lamina propria cells were harvested from the small intestines of K/BxN (black) and *Btk^−/y^*/K/BxN (white) mice as in Methods. Flow cytometry was used to identify the indicated subsets. (A) Representative plots depict B cells in K/BxN (left) and *Btk^−/y^*/K/BxN (right) mice among live lymphocytes. (B) Total B cell frequencies (left) and numbers (right) are shown. (C) Representative plots show IgA^+^ B cells among B220^+^ CD19^+^ live lymphocytes. (D) IgA^+^ B cell frequencies (left) and numbers (right) are shown. (E-G) plasmablasts (surface Igκ^+^), plasma cells (surface Igκ^−^), IgA^+^ plasmablasts (surface Igκ^+^, surface IgA^+^), and IgA^+^ plasma cells (surface Igκ^−^, intracellular IgA^+^) were identified among B220^−^ CD19^−^ intracellular Igκ^+^ CD138^+^ live cells. (E) Representative plots depict the indicated subsets in K/BxN (left) and *Btk^−/y^*/K/BxN (right). (F) Cell subset numbers are shown for plasmablasts (top left), IgA^+^ plasmablasts (top right), plasma cells (bottom left), and IgA^+^ plasma cells (bottom right). n ≥ 6 8-9 wks old mice per group, no significant differences were observed between K/BxN and *Btk^−/y^*/K/BxN groups, two-tailed t test.

### Loss of BTK reduces small intestinal but not serum levels of IgA in K/BxN mice

Whereas intestinal production of IgA is severely compromised in germ-free mice, serum IgA levels are only reduced by half, suggesting that the microbiota contributes to IgA production in the gut whereas a different process is driving IgA production that enters circulation in the serum ^6^. Although IgA levels in *Btk*-deficient mice have long been known to be normal ^14^, we measured IgA in small intestinal lavage, and feces as well as serum of *Btk^−/y^* K/BxN mice. As expected, serum concentrations of IgA were not significantly different between *Btk^−/y^* vs. *Btk^+/y^* K/BxN mice (Fig. 4A). In contrast, levels of free (non-bacterial-bound) IgA were significantly reduced in small intestinal and fecal lavage harvested from *Btk^−/y^* vs. *Btk^+/y^* K/BxN mice (Fig. 4B-C). Our data are consistent with separate routes of serum versus intestinal IgA production.

**Figure 4.**
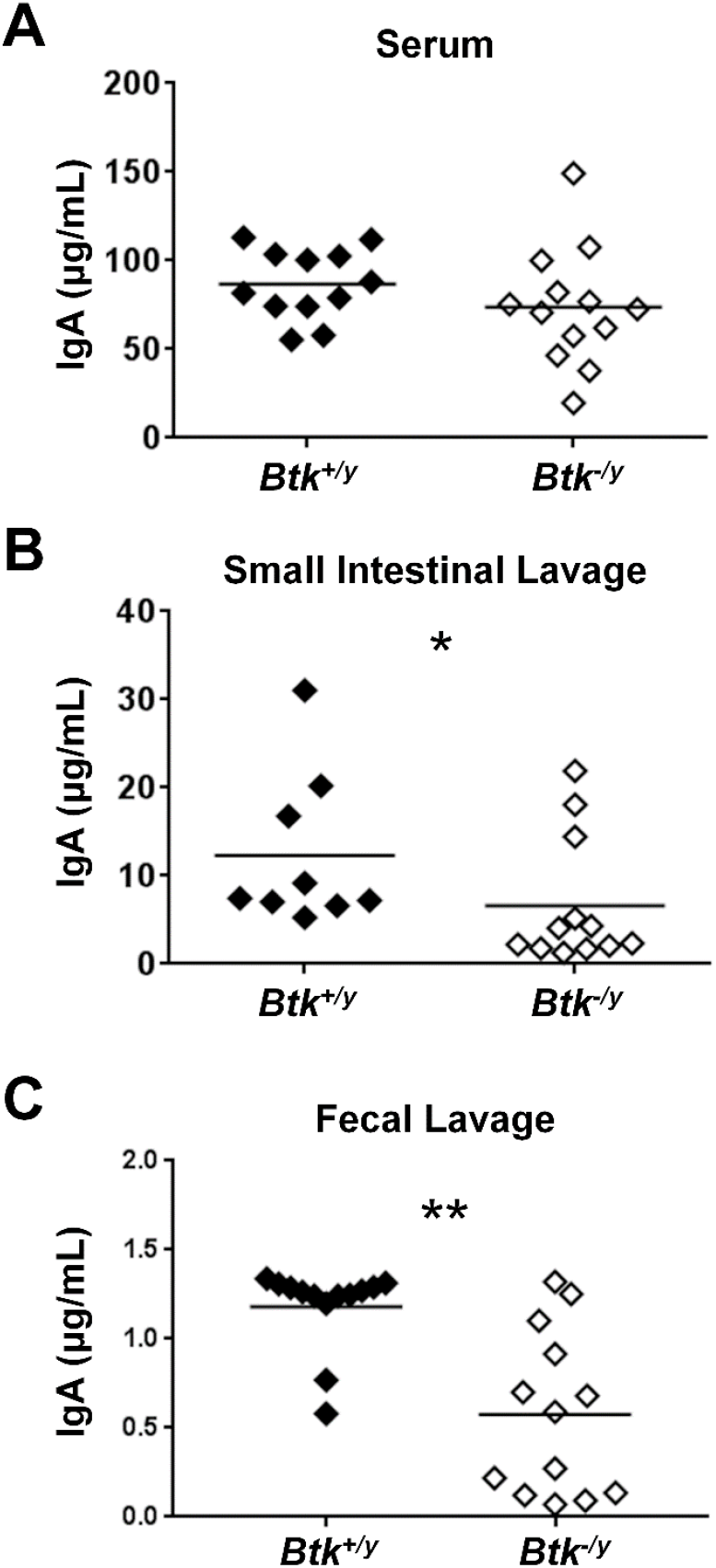
IgA is reduced in the small intestine and feces, but not serum of *Btk*-deficient K/BxN mice. Serum, small intestinal lavage, and fecal lavage samples were collected from 8-9 week-old K/BxN and *Btk^−/y^*/K/BxN mice as in Methods. Total IgA was measured by ELISA as in Methods for A) serum, (B) small intestinal lavage, and (C) fecal lavage. n ≥ 9 mice per group, * p < 0.05, ** p < 0.01, Mann-Whitney U test.

### Disrupted BTK signaling alters CDR3 polarity, but not somatic hypermutation or clonal diversity of intestinal IgA^+^ plasma cells

Intestinal IgA can derive from both T-independent and T-dependent germinal center responses ^5,6^. Given the marked reduction in Peyer’s patch germinal center B cell numbers (Fig. 1) and reduction in free IgA present in the small intestinal lumen of *Btk^−/y^* K/BxN (Fig. 4), we hypothesized the immunoglobulin repertoire of lamina propria IgA plasma cells would be altered. To test this hypothesis, lamina propria cells were isolated from the small intestines of *Btk^+/y^* K/BxN and *Btk^−/y^* K/BxN mice and flow cytometry was used to purify IgA^+^ plasma cells identified as in Fig. 3 (B220^−^ CD19^−^ CD138^+^ intracellular IgA^+^ surface IgA^−^). DNA was extracted and IgH sequencing was performed to assess immunoglobulin gene segment usage, identify somatic mutation rates, and compare amino acid composition between groups. Alpha diversity did not differ significantly between *Btk^+/y^* and *Btk^−/y^* K/BxN IgA plasma cell repertoires (normalized Shannon-Wiener Index, p = 0.82, Mann-Whitney U test). Lamina propria IgA^+^ plasma cells isolated from *Btk^−/y^* versus *Btk^+/y^* K/BxN mice showed no differences in the average rates of mutation in either framework region 3 (FW3) or the V or J regions of complementarity determining region 3 (CDR3) between genotypes (Fig. 5A). A lower rate of mutation was observed in FW3 relative to CDR3, as expected in both groups (p < 0.01 in both genotypes, Mann-Whitney U test). The percentage of mutations resulting in amino acid replacement was not different (not shown). Amino acid properties were also compared between the two genotypes. No difference was observed in CDR3 amino acid charge, disorder, or pH (not shown), but a significant decrease in CDR3 amino acid polarity was observed in IgA^+^ plasma cells isolated from *Btk*^−/y^ K/BxN mice relative to *Btk^+/y^* mice (average: 6.80 vs. 7.28, p = 0.03, Fig. 5B). A trend towards reduced CDR3 length in *Btk*-^−/y^ K/BxN mice was observed but was not significant (p = 0.48, Mann-Whitney U test). These data indicate that *Btk*-^−/y^ IgA^+^ plasma cells undergo somatic hypermutation but show differences in CDR3 composition, suggesting the small intestinal plasma cell repertoire is qualitatively different when BCR signaling is altered.

**Figure 5.**
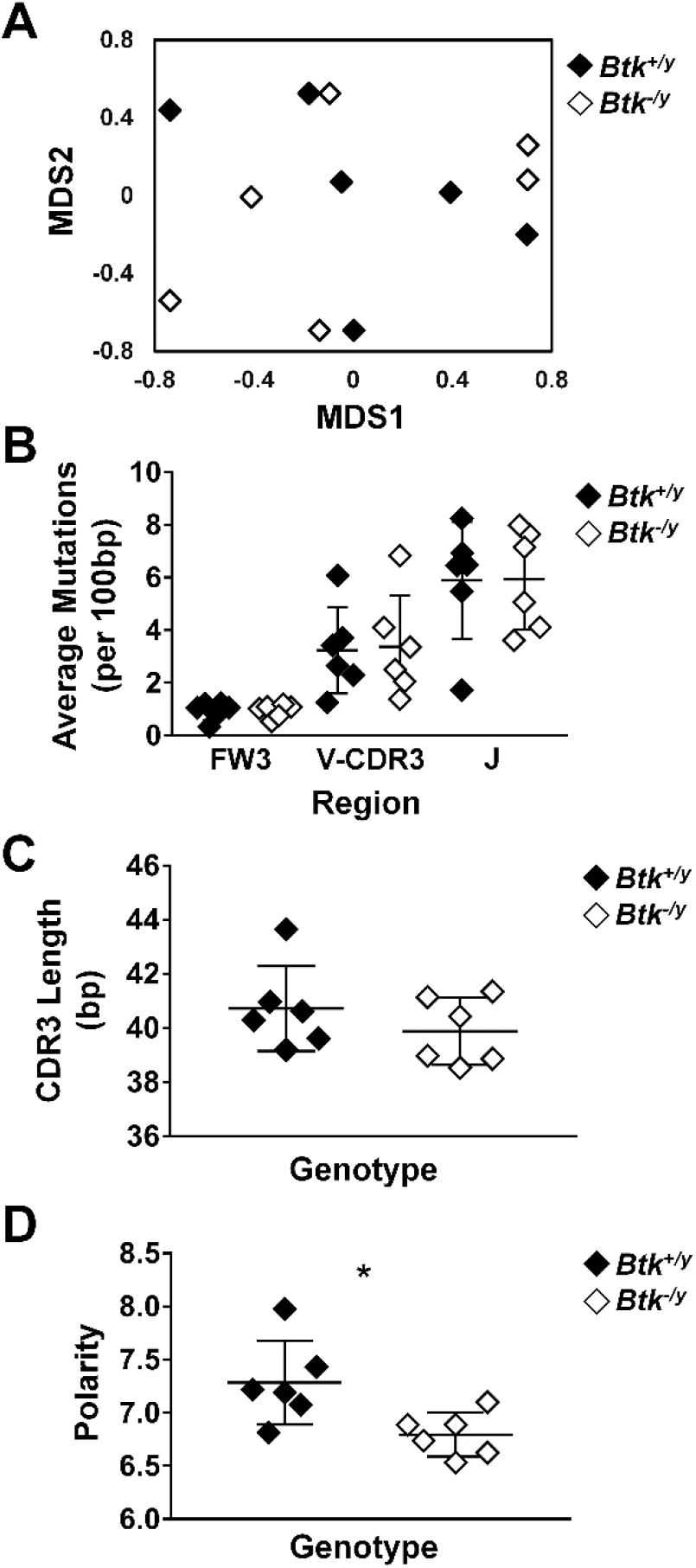
Disrupted BTK signaling does not reduce somatic hypermutation but leads to altered CDR3 polarity in IgA+ plasma cells in the small intestinal lamina propria. Lamina propria cells were harvested from the small intestines of K/BxN (black) and *Btk^−/y^*/K/BxN (white) littermates from 3 litters as in Methods, n = 6 mice per group. IgA^+^ plasma cells were identified as live singlet B220^−^ CD19^−^ CD138^+^, surface IgA^−^, intracellular IgA^+^ cells and flow cytometry sorted. Cells were pelleted, DNA was isolated, and IgH sequencing was performed. An average of 138 productively rearranged sequences was generated per mouse (range: 25-350). Sequence analysis was performed as detailed in Methods. IMGT/HighV-QUEST (http://www.imgt.org/IMGTindex/IMGTHighV-QUEST.php) was used to assign V(D)J genes and assess somatic mutations. VJDtools was used to analyze immunoglobulin gene attributes (https://vdjtools-doc.readthedocs.io/en/master/intro.html). (A) MDS ordination was generated using the Jaccard Index by VDJtools and individual mice are plotted. (B) Somatic mutations in framework (FW) and complementarity determining regions (CDR) of productively rearranged genes and the average number of mutations per 100 bp was calculated for each mouse and is plotted. No significant differences were observed between genotypes, Mann-Whitney U test. (C-D) VDJtools was used to assess CDR3 length (C) and amino acid polarity (D), * p < 0.05, Mann-Whitney U test.

### Disruption of normal B cell signaling decreases IgA coating of commensal bacteria in the small intestine

The small intestine is the major site of interface between B lymphocytes and commensal microorganisms in the gut. To test whether reduced germinal center B cell numbers in Peyer’s patches, peritoneal B1a B cells, and IgA levels in the small intestine of *Btk*-^−/y^ K/BxN mice along with qualitative differences in the IgA^+^ plasma cell repertoire impacted IgA-coating of commensals, bacteria were collected from small intestinal lavage and the frequency of bacteria coated with IgA was assessed using flow cytometry as in Methods. Consistent with previous studies ^12^, *Btk^−/y^* K/BxN mice showed less severe arthritis as compared to *Btk^+/y^* K/BxN mice (average arthritis scores were 4.7 vs. 12.1, respectively, p < 0.001, Mann-Whitney U test, Fig. 6A). The average proportion of IgA-coated bacteria was reduced in *Btk^−/y^* vs. *Btk^+/y^* K/BxN mice (Fig. 6B-C, 44 % vs. 65 %, respectively, p < 0.05). These data show that *Btk* loss impairs IgA-coating of commensal microbes.

**Figure 6.**
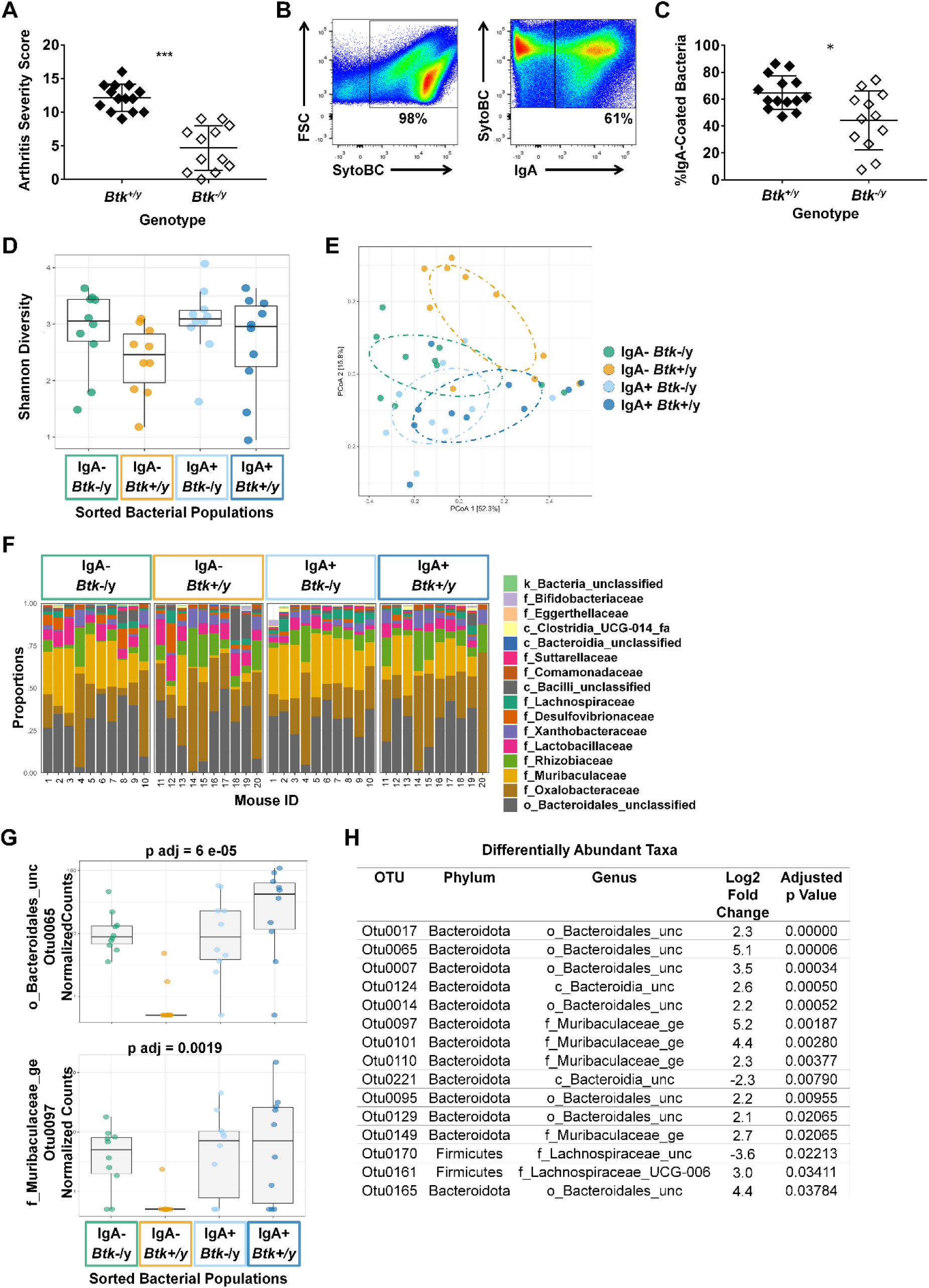
IgA coating of microbes differs between *Btk-*deficient K/BxN mice and *Btk*-sufficient littermates. (A) An arthritis severity score was assigned to each paw (0-4) and the total arthritis severity score was calculated (0-16) for n ≥ 12 *Btk^+/y^* (black) or *Btk^−/y^* (white) K/BxN mice per group at the time of small intestinal harvest, p < 0.001, Mann-Whitney U test. (B-H) Bacteria were harvested from the small intestinal lavage of K/BxN and *Btk^−/y^*/K/BxN littermates from 5 litters that were 8-9 wks old. Cells were stained with SytoBC and anti-IgA antibody and flow cytometry was used to identify and sort IgA-coated or uncoated bacteria. (B) Representative plots. (C) The frequency of IgA-coated bacteria is plotted for n ≥ 12 individual *Btk^+/y^* and *Btk^−/y^* K/BxN mice per group, p < 0.05, Mann-Whitney U test. (D-H) DNA was isolated from bacteria sorted from *Btk^+/y^* and *Btk^−/y^* K/BxN mice based on being IgA coated or uncoated and microbiome sequencing and analysis was performed by Microbiome Insights using Mothur for the following four groups: *Btk^−/y^* IgA- (green), *Btk^+/y^* IgA- (gold), *Btk^−/y^* IgA+ (light blue), *Btk^+/y^* IgA+ (dark blue). The average number of quality filtered sequence reads per sample was 35446 (Range: 10,000-50,000). (D) Shannon diversity for each mouse is plotted; *Btk*: p = 0.06, IgA: p = 0.33, IgA:Btk interaction: p=0.81, ANOVA. (E) Individual mice are plotted based on principle component analysis (PCoA). (F) Proportions of the indicated taxa at the family level are plotted for each of n = 10 mice per group. (G-H) The DESeq2 package was used to identify differentially abundant taxa among IgA variables using a linear model that included IgA, BTK, and BTK*IgA interactions. Normalized counts for two OTUs and corresponding adjusted p values are shown in panel G; other differentially abundant OTUs are shown in panel H and Fig. S1).

### Btk-deficient mice can produce IgA in response to commensal microbes, but some bacteria escape into the IgA-free fraction

To assess whether loss of Btk altered the community composition between IgA-coated and uncoated bacteria in the small intestine, bacterial community composition was compared using the previously described IgA-seq technique ^22,23^. IgA coated versus uncoated microbes were flow cytometry-sorted, 16S sequenced, and analyzed (Fig 6B-H). No significant differences in Shannon (alpha) diversity were observed, although diversity trended higher in *Btk^−/y^* K/BxN (Fig. 6D). Microbial composition was compared across the four groups. OTU abundances were summarized into Bray-Curtis dissimilarities and a PCoA ordination was performed to obtain graphical representation of microbiome composition similarity among samples: IgA^−^/*Btk^−/y^* (green), IgA^−^/*Btk^+/y^* (gold), IgA^+^/*Btk^−/y^* (light blue), IgA^+^/*Btk^+/y^* (dark blue) (Fig 6E).

Permutational analysis of variance (PERMANOVA) determined significant differences in beta-diversity among IgA (p=0.014), and BTK (p=0.013) factors. One group appeared to stand out from the others: uncoated bacteria from mice with normal B cells, or IgA^−^/*Btk^+/y^* (gold). This was expected, as IgA-uncoated bacteria are known to differ from IgA-coated bacteria ^22,23^, and indeed, IgA coated versus uncoated bacteria in these mice with normal B cells separated well in the PCoA plot (IgA^−^/*Btk^+/y^* (gold) vs IgA^+^/*Btk^+/y^* (dark blue) and trended toward significance in pairwise analysis (p=0.08). This contrasts IgA coated and uncoated commensals from *Btk*-deficient mice (IgA^−^/*Btk^−/y^* (green) vs IgA^+^/*Btk^−/y^* (light blue), which overlap more in the PCoA and are not statistically different by pairwise analysis (p=0.14). Interestingly, there is near-complete overlap between IgA coated bacteria from *Btk*-deficient and -sufficient mice (IgA^+^/*Btk^−/y^* (light blue), IgA^+^/*Btk^+/y^* (dark blue, p=0.13), indicating that the *Btk*-deficient B cells can respond to the same microbes that elicit IgA in normal B cells.

A per sample view of taxonomic composition showing the most abundant taxa at the family level is shown in Fig 6F. Each bar represents a single sample, and they are grouped as in the PCoA plot. Again, the IgA^+^ bar graphs appear similar between *Btk*-deficient and sufficient mice, and both of these look similar to the IgA-uncoated fraction from *Btk*-deficient mice. In contrast, IgA-uncoated bacteria from *Btk*-sufficient mice (normal B cells) again differ from the other three, most prominently in lacking *Muribaculaceae* (Fig. 6H). At the genus level, 15 differentially abundant taxa were significantly different when tested for the interaction between *Btk* genotype and IgA coating (Figs 6H and S1). Twelve of these show a distinct pattern in which the OTU is highly abundant in IgA^+^ fractions from both genotypes, as well as the IgA-negative fraction in the *Btk*-deficient samples, but is not abundant in the IgA-negative fraction of the *Btk*^+/y^ samples (Figs 6G and S1). Again, *Muribaculaceae* are well-represented among these, as are *Bacteroidales*. Overall, these findings indicate that IgA from *Btk*-deficient B cells can bind the same commensal bacteria as IgA from normal B cells, but is deficient, either in abundance or affinity, resulting in inadequate coating, and escape of some microbes into the IgA uncoated fraction.

### Parabacteroides distasonis is reduced in autoimmune arthritis-protected Btk^−/y^ K/BxN mice

Feces sampling is much more practical for clinical applications and for longitudinal studies. Feces primarily reflect the large intestinal microbiome, which differs markedly from the small intestinal community ^24^. We therefore assessed microbiome differences present in the feces of *Btk^+/y^* and *Btk^−/y^* K/BxN mice. Large intestinal alpha and beta diversity were not different between *Btk*-deficient and *Btk*-sufficient K/BxN mice (p = 0.96, Kruskal-Wallis and p = 0.63, PERMANOVA, respectively, Fig. 7). However, one specific OTU was identified as significantly increased in *Btk*-sufficient K/BxN. This OTU was identified down to the species level as *Parabacteroides distasonis* (p <0.05). *P. distasonis* is a gram-negative obligate anaerobe that belongs to the *Porphyromonadaceae* family ^25^.

**Figure 7.**
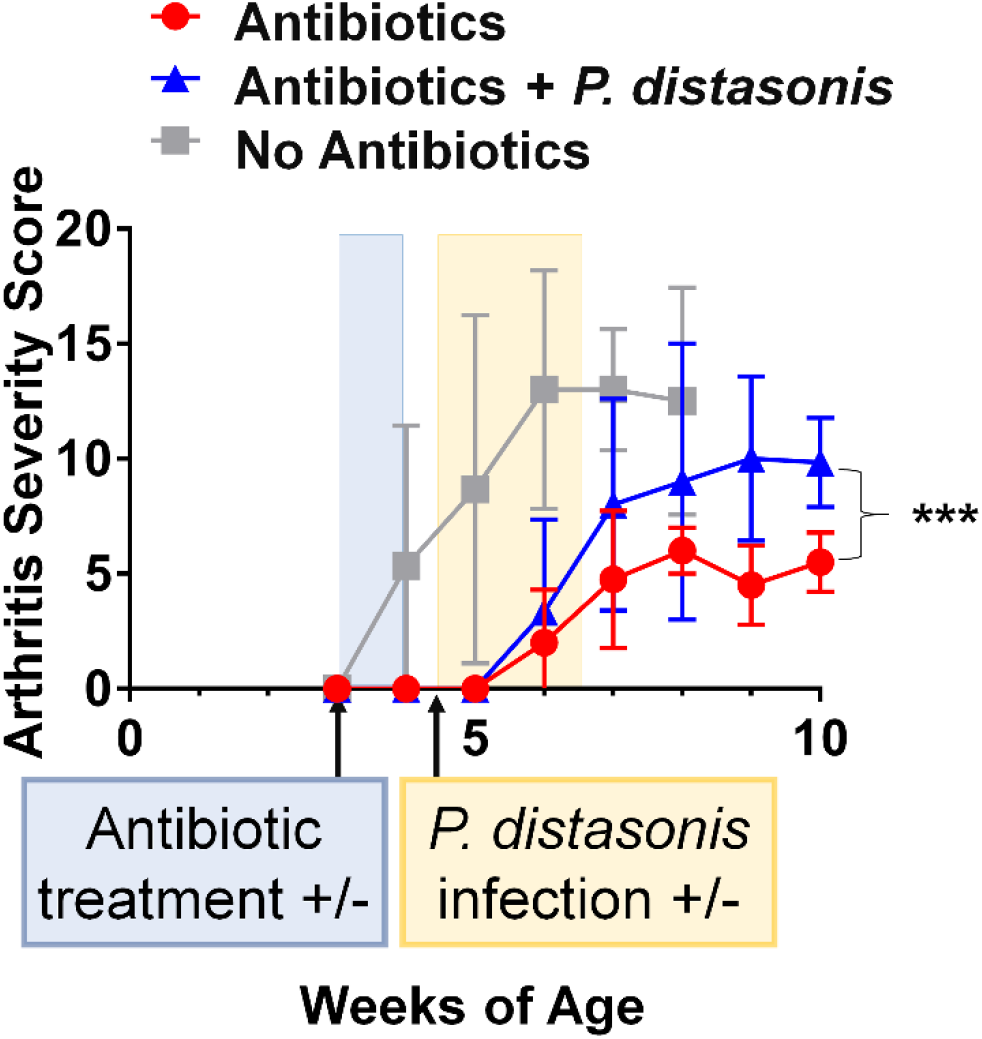
*Parabacteroides distasonis* increases arthritis severity in K/BxN mice that are otherwise protected by prior antibiotic treatment. K/BxN mice were treated at weaning with antibiotics for 1 week (pale orange box), followed by oral gavage with saline or 4.0×10^7^-1.5×10^8^ CFU of *Parabacteroides distasonis* (pale blue box) per mouse three times per week for two weeks. Arthritis severity was scored weekly for K/BxN mice that received no antibiotics (grey squares, n = 3), antibiotics (red circles, n = 4), or antibiotics followed by *P. distasonis* infection (blue triangles, n = 6), *** p < 0.001, two-way ANOVA.

### Parabacteroides distasonis drives autoimmune arthritis in antibiotic-protected K/BxN mice

*P. distasonis* was increased in the feces of arthritis-prone K/BxN mice relative to protected *Btk^−/y^* K/BxN mice, suggesting these commensals may play an inciting role in autoimmune arthritis. To answer this question, we chose to treat K/BxN mice with antibiotics at weaning, as this reduces autoimmune arthritis ^21^ and provides the opportunity to test the impact of infection with a commensal of interest. Importantly, this model avoids the caveat of disrupted gut immune system and structure development observed in germ-free mice. K/BxN mice were therefore administered antibiotics at weaning for 1 week, followed by 2 days to allow antibiotic washout, and were subsequently infected with *P. distasonis* by oral gavage three times per week for two weeks. One group of K/BxN control mice received antibiotics followed by oral gavage of saline as a placebo, and another received no treatment of any kind. Arthritis severity was monitored weekly until 8-10 weeks of age. Control mice that received no antibiotics or gavage developed autoimmune arthritis, as expected, and were euthanized at 8 weeks of age due to arthritis severity (Fig. 7D). In comparison, mice treated with antibiotics and placebo gavage were protected from arthritis (p < 0.0001, two-way ANOVA). However, reconstitution of antibiotic-treated mice with *P. distasonis* resulted in a significant increase in arthritis scores compared to those that got only placebo (p < 0.0001, two-way ANOVA). These data suggest that *P. distasonis* is capable of enhancing autoimmune arthritis in mice that are protected from disease by microbiome suppression.

## Discussion

The microbiome, shaped by IgA, affects autoimmunity in complex ways ^7,21,22,26–29^. However, until now the contributions of B lymphocyte signaling to these processes had not been studied. The work presented here uses the *Btk*-deficient K/BxN mouse model to reveal that mucosal germinal center B cells and intestinal IgA rely in part on this B cell signaling protein (Figs 1, 4). In the absence of *Btk*, gut IgA is reduced, and is impaired in its ability to coat bacteria (Figs 4, 6). Interestingly, even in this suboptimal state, *Btk*-deficient B cells can still produce bacteria-specific IgA (Fig 6) and generate enough IgA^+^ plasma cells to populate the lamina propria (Fig. 3). However, this IgA is functionally incomplete, since some microbes that should be IgA-coated are found to be IgA-free, particularly those in the *Muribaculaceae* and *Bacteroidiales* families (Fig. 6). While the fecal microbiome in this model does not appear to be substantially shifted, one organism, *Parabacteroides distasonis,* may be disadvantaged in the setting of *Btk*-deficiency, as it is found in higher abundance in *Btk*^+^ K/BxN. The reason for this may be defective IgA responses in *Btk*-deficient mice, as some bacteria rely on normal IgA for signaling activation or for adherence to the mucous layer ^9,10^. Alternatively, the species that are able to escape IgA binding in *Btk*-deficient mice may then be able to outcompete *P. distasonis.* The ability of *P. distasonis* to trigger arthritis when reconstituted into antibiotic-treated, disease-protected, K/BxN mice (Fig 7) suggests that its loss in *Btk*-deficient mice could be a factor in disease-protection.

The source of the IgA that remains in *Btk*-deficient mice requires further study. IgA^+^ plasma cells are present in normal numbers in the lamina propria (Fig. 3), despite greatly reduced numbers of the B cells that give rise to them. IgA^+^ plasma cells normally arise from innate-like B1 as well as standard B2 B cells. 25-50% of intestinal IgA produced in mice is B1 B cell-derived and the remaining 50-75% is produced by B cells from the B2 lineage ^5,6^. As in non-autoimmune strains ^14^, B1 cells are nearly absent from *Btk*-deficient K/BxN mice (Fig. 2), making their contribution to the population of intestinal plasma cells unlikely. B2 cells normally form very large, active GCs in PPs. The reduced numbers of GC B cells in PPs are consistent with previous work demonstrating equally dramatic reduction of GC B cells in joint-draining popliteal lymph nodes of *Btk*-deficient K/BxN mice ^12^, and imply a general role for BTK in GC development or function. Interestingly however, while numbers are reduced, frequencies of IgA^+^ and GC B cells in the Peyer’s patches do not differ, suggesting that *Btk*-deficient B cells are still capable of IgA class-switching despite impaired BCR signaling. This is also consistent with *Btk*-deficient T-dependent IgG responses that, while blunted at primary challenge, are adequate following subsequent T-dependent boost. ^14–16,18,30^. Preserved T-dependent responses to commensals may similarly support the small reservoir of GC B cells in PPs of *Btk*-deficient K/BxN mice. In addition, the extent of mutation observed in CDR3 (~5 mutations per 100bp, Fig. 5) suggests B2 origin of IgA^+^ plasma cells, rather than B1, which tend to have lower levels of somatic hypermutation ^31^. Our findings are also in line with previous data showing that somatic hypermutation still occurs in Peyer’s patch GC B cells even when surface BCR expression (and thus BCR signaling) is ablated ^32^. Overall, the data suggest that B2 B cells in *Btk*-deficient K/BxN mice are able to undergo T-dependent interactions, and it seems likely that the few precursors are able to expand homeostatically to generate adequate plasma cells to populate the lamina propria. Further study of full length BCR sequences is needed to elucidate clonal relationships and better define the range of somatic hypermutation in these cells.

In non-autoimmune mice, free IgA in the gut is primarily derived from B2 cells, whereas bacterial-bound IgA derives from both B1 and B2-B cell lineages ^22^. Despite normal numbers of IgA^+^ plasma cells in the lamina propria, free IgA present in the small and large intestine was reduced in *Btk*-deficient K/BxN mice (Fig. 4). Two possible interpretations are 1) IgA production on a per cell level is reduced or 2) IgA production is normal but a higher proportion of IgA is bound to commensal microbes, leaving less free IgA for detection in *Btk*-deficient mice. Our data suggest the first possibility, as the frequency of IgA-coated bacteria is *lower* in *Btk*-deficient K/BxN mice (Fig. 6C). In support of this interpretation, a BTK/Ets1/Blimp-1 axis is implicated in per-cell antibody secretion as follows. Reduced steady state serum levels of IgM and IgG in *Btk*-deficient mice is normalized in *Btk*-deficient/*Ets1*-deficient mice ^33^. Ets1 is a transcription factor that physically interacts with Blimp-1 to limit its function ^34^. Blimp-1 elimination permits plasma cell survival but limits the amount of antibody secreted per cell ^35^. Thus, *Btk*-deficiency leaves Ets1 activity unopposed which results in diminished Blimp-1 activity and, in turn, reduced levels of antibody secretion by plasma cells.

The lower levels of microbial IgA coating from *Btk*-deficient K/BxN initially raised the possibility that commensal-driven IgA production was lacking, perhaps due to impaired GC processes, resulting in failure of the B cells to respond to bacteria that normally elicit IgA. Therefore it was surprising to find that IgA from *Btk*-deficient mice did target the same bacteria as that from normal B cells: there was no difference in IgA-coated community composition between *Btk*-deficient and *Btk*-sufficient K/BxN mice (Fig. 6 E,F). This indicates that BTK is not required to support specific commensal-driven production of IgA, at least in the K/BxN model. However, the affinity for antigen epitopes may be lower, or the quantity of IgA may be insufficient, to fully cover all of the targeted commensals, thus allowing some of them to escape IgA coating, as indicated by the shifting of uncoated IgA samples from *Btk*-deficient mice to more closely resemble IgA-coated fractions. This profile included many OTUs present exclusively in IgA^+^ samples from *Btk*-sufficient animals: 12 individual, differentially-expressed OTUs were almost universally IgA coated in *Btk^+/y^* mice, contrasting their equal abundance in IgA^+^ and IgA^−^ fractions from *Btk*^−/y^ mice (Fig. 6 G,H, S1).

Because small intestinal microbiome samples are difficult to acquire in humans, fecal samples are generally used, and microbiome alterations are associated with many disease states, including rheumatoid arthritis ^36^. Therefore, we also compared fecal alpha and beta diversity between *Btk*-deficient and *Btk*-sufficient K/BxN mice but found no differences. In fact, the only OTU found to be significantly difference in the two groups was *P. distasonis*, which was unexpectedly higher in the setting of *Btk*-sufficiency. *P. distasonis* is a prototype commensal, as it is a member of the altered Schaedler flora (ASF) that is used to colonize germ-free mice with a minimally diverse microbiome ^37^. ASF colonization has been shown to drive production of Th17 and Th1 responses in a transgenic system deficient in Tregs, suggesting that commensals can, under the wrong circumstances, drive T cells to acquire an inflammatory phenotype ^38^. Our data show that *P. distasonis* is increased in association with autoimmune arthritis in *Btk^+/y^*/K/BxN mice, and that it can support autoimmune arthritis development in mice that are otherwise protected by antibiotic elimination of commensals (Fig. 7). This organism is unlikely to provide an explanation for the entire disease process, as it did not provide as robust an arthritis as that found in mice not treated with antibiotics, but may be one factor. Of note, Segmented Filamentous Bacteria (SFB), now known as *Candidata savagella*, has been shown to drive arthritis in this model via TH17 enhancement ^21^, but was not differentially expressed in any of our comparative data. Thus, there are likely a number of bacteria that can trigger arthritis.

Interestingly, *P. distasonis* does not uniformly trigger autoimmunity. It is reduced in the feces of multiple sclerosis patients, where it drives CD4^+^ IL-10^+^ T cell differentiation ^28^. Treatment of germ-free mice with the membranous fraction of *P. distasonis* lysates induces FoxP3+ Treg cells in mesenteric lymph nodes, but not Peyer’s patches, and reduces DSS-induced colitis severity ^29^. Therefore, context of the host must thus be taken into account when considering the impact of commensal organisms on gut immune responses.

One additional consideration for this model is that BTK also plays a role in myeloid cells and mast cells. Although it was previously shown that BTK in myeloid cells was not necessary for arthritis in the K/BxN serum transfer model ^12^, this question has not been tested in the more robust spontaneous K/BxN model. In addition, BTK is also important for TLR4 responses to LPS in addition to BCR responses, which is not independently analyzed here, but could be an important contributor ^14^. Finally, the T cell transgene that drives arthritis in K/BxN mice may also alter the immune response to commensals, including, but not limited to, changes in T cell help to B cells.

Clinical use of BTK inhibitors is likely to continue to increase given the number of disease applications for which they are either approved or are being tested. Recent studies show that BTK is important for the maturation but not survival of autoreactive-prone B cells and that BTK signaling is needed for normal T-independent type II responses in a model in which *Btk* deletion can be timed to mimic the scenario of clinical inhibition ^39^. The data presented here suggest that microbiome perturbation is a likely outcome of BTK inhibition. This should be carefully considered with respect to individual patients, as changes that are beneficial in one setting (e.g. arthritis disease prevention by antibiotic treatment in autoimmune K/BxN mice) can be harmful in another (e.g. type 1 diabetes exacerbation by antibiotic treatment in NOD mice) ^21,40^. Overall, these data show for the first time that BTK-signaling contributes to IgA development and interaction with commensal bacteria and may alter the microbiome in ways that affect autoimmunity.

## Methods

### Animals and disease assessment

*Btk^−/−^*/NOD mice were backcrossed to the NOD strain >20 generations and were generated as described previously ^13^. KRN C57BL/6 mice were a gift from Cristophe Benoist and Diane Mathis (Harvard Medical School, Boston MA). KRN male mice were bred to *Btk*^+/−^/NOD female mice to generate *Btk^+/y^*/K/BxN and *Btk^−/y^*/K/BxN littermates. Mice were scored for arthritis beginning at weaning by the Chondrex mouse arthritis scoring system (https://www.chondrex.com/documents/Mouse%20CIA.pdf). Paw thickness of the hind footpad was measured using a dial gauge (13-159-9, Swiss Precision Instruments). All mice were housed under specific pathogen-free conditions, or in the case of *P. distasonis* infection, in the ABSL-2 facility, and all studies were approved by the institutional animal care and use committee of Vanderbilt University, fully accredited by the AAALAC.

### Mammalian cell isolation, flow cytometry, and antibodies

Small intestines were dissected and intestinal contents were flushed by lavage with 5mL 1X PBS + EDTA-free Complete Mini Protease Inhibitor Cocktail Tabs (Roche). Peyer’s patches were dissected from the small intestine and macerated in HBSS + 10% FBS. Lamina propria cells were isolated after Peyer’s patches were removed as follows. Intestines were cut longitudinally and mucus was gently scraped off and discarded. Tissue was cut into ~0.5 cm pieces and incubated with HBSS + 5% BCS + 2 mM EDTA at 37°C for 45 min while shaking. Recovered tissue was washed, minced, and incubated with 1.5 mg/ml collagenase VIII (from *Clostridium histolyticum*, Sigma-Aldrich) and 100 U DNase I (Sigma-Aldrich) at 37°C for 25 min while shaking. Cells were vortexed for 20 s and HBSS + 5% BCS was added, cells were pelleted, and resuspended in 40% Percoll (GE Healthcare). 70% Percoll was underlayed and the interface layer was recovered after centrifugation at 300 x g for 20min at 4°C. Cells were pelleted, resuspended, and counted.

Cells were stained for flow cytometry in 1X PBS containing 0.1% sodium azide, 0.02% EDTA, and 0.05% FBS using the following reagents and reactive antibodies: B220 (6B2), CD19 (1D3), CD95/FAS (JO2), CD138 (281-2), GL7, Igκ (187.1), or Ghost Dye (BD Biosciences, eBioscience, or Tonbo Biosciences), or IgA (1040-09, 1040-02, or 1040-31, Southern Biotech). Lamina propria cells were stained intracellularly for CD138, as this protein was cleaved from the cell surface during the isolation protocol (not shown). The Cytofix/Cytoperm kit was used for intracellular permeabilization and staining (BD Biosciences). Samples were acquired using a BD Biosciences LSR II flow cytometer and FlowJo software (Tree Star, Inc) was used for data analysis.

### Fecal and small intestinal lavage sample preparation and ELISA

Freshly collected feces were resuspended at 0.1mg/mL in 1X PBS containing EDTA-free Complete Mini Protease Inhibitor Cocktail Tabs (Roche) per the manufacturer’s instructions. Samples were subsequently spun at 4000 RPM for 10 min., supernatant was removed and spun for an additional 10min and resulting supernatant was sterile filtered through 0.22um filters and stored at −20°C.

For detection of total IgA, plates were coated with goat anti-mouse IgM + IgG + IgA (H+L) (SouthernBiotech) diluted in 1X PBS, followed by blocking with 10% non-fat dry milk. Sera were diluted 1:50,000 in 1X PBS + 0.05% tween and incubated 2h at room temperature. Bound IgA or IgM antibody was detected by incubation with anti-IgA-alkaline phosphatase (goat anti-mouse IgA (α chain-specific) (SouthernBiotech) or anti-IgM-alkaline phosphatase goat anti-mouse IgM (μ chain-specific) (SouthernBiotech), respectively diluted in veronal buffered saline (142 mM NaCL + 5 mM sodium barbital) + 15mM sodium azide + 0.5% FBS. Plates were washed 4X with 1X PBS + 0.05% tween after each incubation step. 10mg/mL *p*-nitrophenyl phosphate substrate (Sigma-Aldrich) was added and O.D. was read at 405 nm *using a* Microplate Autoreader (Bio-Tek Instruments). Intestinal IgA was quantified based on a standard curve that was generated using Purified Mouse IgA, κ Isotype Control (BD Biosciences).

### Plasma cell IgH sequencing

Lamina propria cells prepared from freshly isolated small intestines as outlined above. Cells were stained for surface proteins, fixed with 0.5% paraformaldehyde for 5 minutes (to retain surface antibodies but limit potential DNA cross-linking), washed, and permeabilized with 0.1% saponin in 1X PBS containing 10% bovine calf serum, followed by staining for intracellular proteins using antibody clones outlined above. A BD FACSAria III was used to sort IgA^+^ plasma cells that were resuspended in 1X PBS (live B220^−^ CD19^−^ intracellular IgA^+^ surface IgA^−^ CD138^+^ cells). Cell pellets were stored at −80°C prior to DNA extraction. DNA was isolated as a single batch using the QIAamp DNA Micro Kit (Qiagen) per the manufacturer’s instructions. Mouse IgH CDR3 PCR amplification and sequencing was performed using the ImmunoSeq assay and sequence data were demultiplexed/filtered/processed by Adaptive Biotechnologies (Seattle, WA) ^41^. IMGT/HighV-QUEST (https://www.imgt.org/HighV-QUEST/) was used to assign gene identities and identify mutations among sequences with productive rearrangements. VDJtools (https://vdjtools-doc.readthedocs.io/en/master/index.html) was used to compare CDR3 properties between the two groups.

### Microbiome sample preparation and sequence analysis

For IgA-seq studies, small intestines were dissected and intestinal contents were flushed by lavage as above and centrifuged at 400 x g for 5 min. Supernatant was then centrifuged at 8000 x g for 10 min to pellet bacteria. The bacterial pellet was resuspended in 1X PBS + 0.25% BSA + 5% NGS and stained with Fc Block (BD Biosciences, clone 2.4G2), followed by SYTO BC (ThermoFisher) and goat anti-mouse IgA. Cells were sorted on a BD FACSAria III and cells were pelleted and stored at −80°C. DNA extraction, microbiome sequencing, and analysis was performed as a single batch by Microbiome Insights. Bacterial 16S rRNA v4 regions were PCR-amplified as in ^42^. Amplicons were sequenced with an Illumina MiSeq using the 250-bp paired-end kit (v.2). Sequences were denoised, taxonomically classified (Greengenes v. 13_8 reference database), and clustered into 97%-similarity operational taxonomic units (OTUs) using Mothur, v. 1.39.5 ^43^. Potential contamination was addressed by co-sequencing DNA amplified from specimens and from n=4 template-free controls and n=4 extraction kit reagents processed the same way as the specimens. Two positive controls, consisting of cloned SUP05 DNA, were also included (number of copies = 2*10^6). OTUs were considered putative contaminants (and were removed) if their mean abundance in controls reached or exceeded 25% of their mean abundance in specimens. Fecal microbiome analysis is outlined in Supplemental Data. Microbiome statistical analysis approaches are detailed in figure legends.

### Bacterial culture and murine infection

*Parabacteroides distasonis* (ATCC 8503) was cultured in MTGE Anaerobic Enrichment broth (Anaerobe Systems) in an anaerobic chamber. Colony forming unit (CFU) assays were performed to enumerate the number of bacteria used for infection studies. Mice were administered 1mg/mL ampicillin, 1mg/mL neomycin, 1 mg/mL metronidazole, and 0.5 mg/mL vancomycin in drinking water including 2.5 mg/mL Equal sweetener at weaning (3 wks of age) for 7 days to reduce commensal microbes. Mice were then transferred to autoclaved drinking water for 2d and then infected with 4.0×10^7^-1.5×10^8^ CFU *P. distasonis* resuspended in 1X PBS by oral gavage 3 times weekly for 2 wks. All antibiotic drugs were purchased from VWR/Amresco.

## Author contributions

R.H.B., C.E.T., L.E.N., and B.B.B. performed experiments. R.H.B., C.E.T., L.E.N., and C.S.W. performed data analysis. R.H.B. wrote the manuscript, which P.L.K critically reviewed. All authors provided helpful experimental discussion.

## Data availability

The raw sequence reads for all microbiome studies described here are deposited in the NCBI Sequence Read Archive (SRA) Database, BioProject ID: PRJNA706065. BCR repertoire data are available through ImmuneACCESS, https://clients.adaptivebiotech.com/pub/bonami-2021-biorxiv (Adaptive Biotechnologies, Seattle, WA).

## Acknowledgements

Flow cytometry experiments were performed in the Vanderbilt Medical Center Flow Cytometry Shared Resource. Immunoglobulin heavy chain sequencing was performed by Adaptive Biotechnologies (Seattle, WA). We would like to thank Wyatt McDonnell (Vanderbilt University, Nashville, TN) for immunoglobulin analysis advice. Microbiome sequencing and analysis was performed by Microbiome Insights (Vancouver, BC) and Second Genome (San Francisco, CA). We would like to thank Dr. Egle Cekanaviciute (University of California, San Francisco, CA) for providing *Parabacteroides distasonis* culture and infection protocols.

