## Supplemental Methods for "Bruton’s Tyrosine Kinase Supports Gut Mucosal Immunity and Commensal Microbiome Recognition in Autoimmune Arthritis"

*Fecal microbiome analysis*

Fecal pellets were collected and frozen at -20^o^C. DNA extraction, V4 region PCR amplification, sequencing, and statistical analysis was performed as a single batch by Second Genome using Mothur. Amplicons were sequenced with an Illumina MiSeq, sequences were denoised, taxonomically classified (Greengenes reference database), and clustered into 97%-similarity operational taxonomic units (OTUs) using Mothur (42).
