## Supplemental Figure 1 for "Bruton’s Tyrosine Kinase Supports Gut Mucosal Immunity and Commensal Microbiome Recognition in Autoimmune Arthritis"

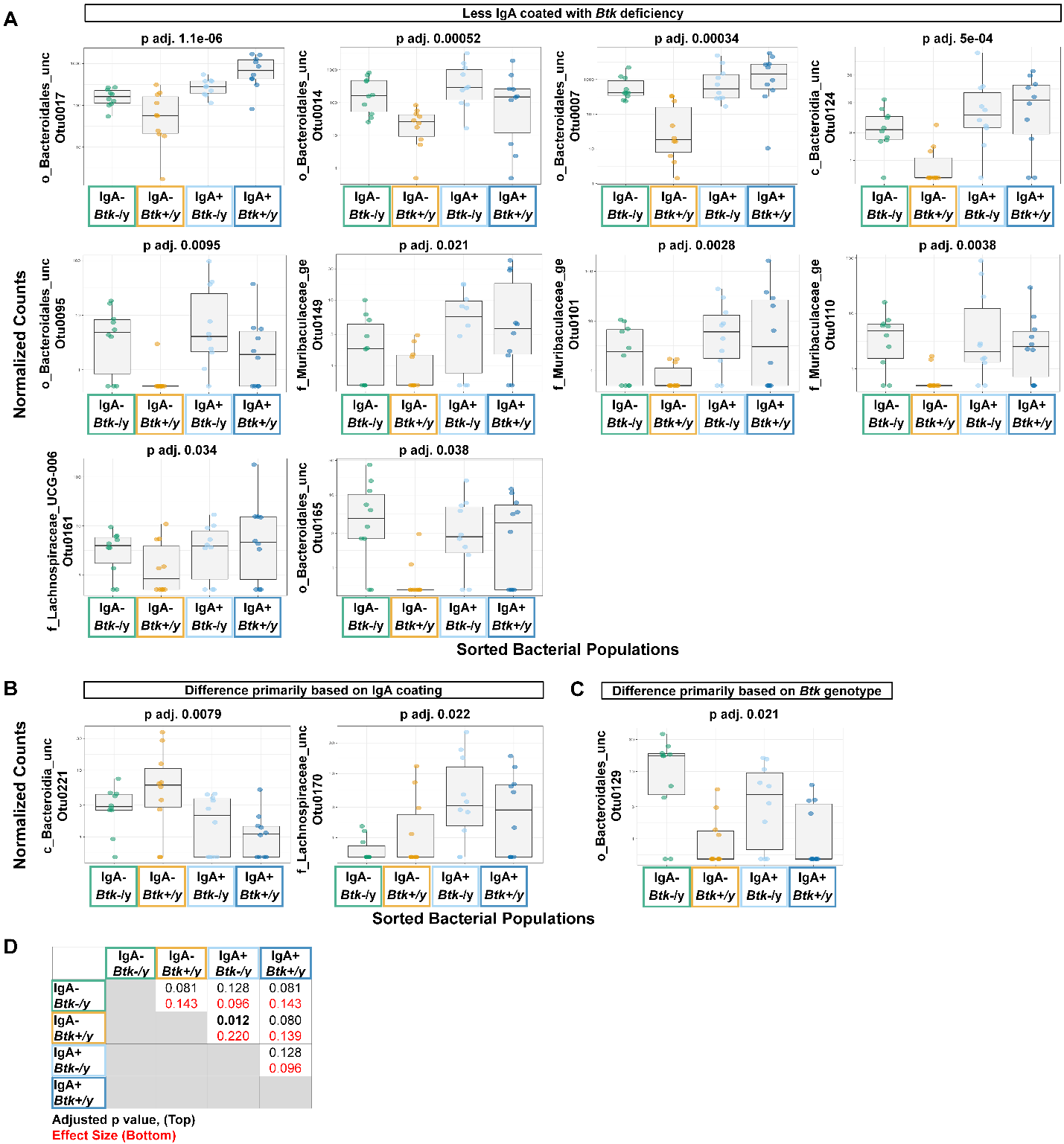

**Supplemental Figure 1. Differentially-abundant bacterial OTUs are identified by IgA-seq analysis of IgA-coated and uncoated commensals from the small intestine of *Btk-*deficient K/BxN mice and *Btk*-sufficient littermates.** Bacteria were harvested from the small intestinal lavage of K/BxN and *Btk^-/y^*/K/BxN littermates (n ≥ 12 individual mice per group) and IgA-seq analysis was performed by Microbiome Insights using Mothur as in Fig. 6 for the following four groups: *Btk^-/y^* IgA- (green), *Btk^+/y^* IgA- (gold), *Btk^-/y^* IgA+ (light blue), *Btk^+/y^* IgA+ (dark blue). The DESeq2 package was used to identify differentially abundant taxa among IgA variables using a linear model that included IgA, BTK, and BTK*IgA interactions. Fifteen differentially abundant OTUs were identified; normalized counts for two of these are shown in Fig. 6 and the remaining thirteen OTUs are shown in panels A-C. Adjusted p values are indicated above each plot, in which individual mice are plotted. These differentially expressed OTUs were grouped as follows: A) Less IgA-coated with *Btk*-deficiency, (B) Difference primarily based on IgA coating, and (C) Difference primarily based on *Btk* genotype. (D) The adjusted p value (black, top) is shown for each pairwise comparison of the four groups using a PERMANOVA (adonis test) along with the effect size (red, bottom).
