## Supplemental Figure 2 for "Bruton’s Tyrosine Kinase Supports Gut Mucosal Immunity and Commensal Microbiome Recognition in Autoimmune Arthritis"

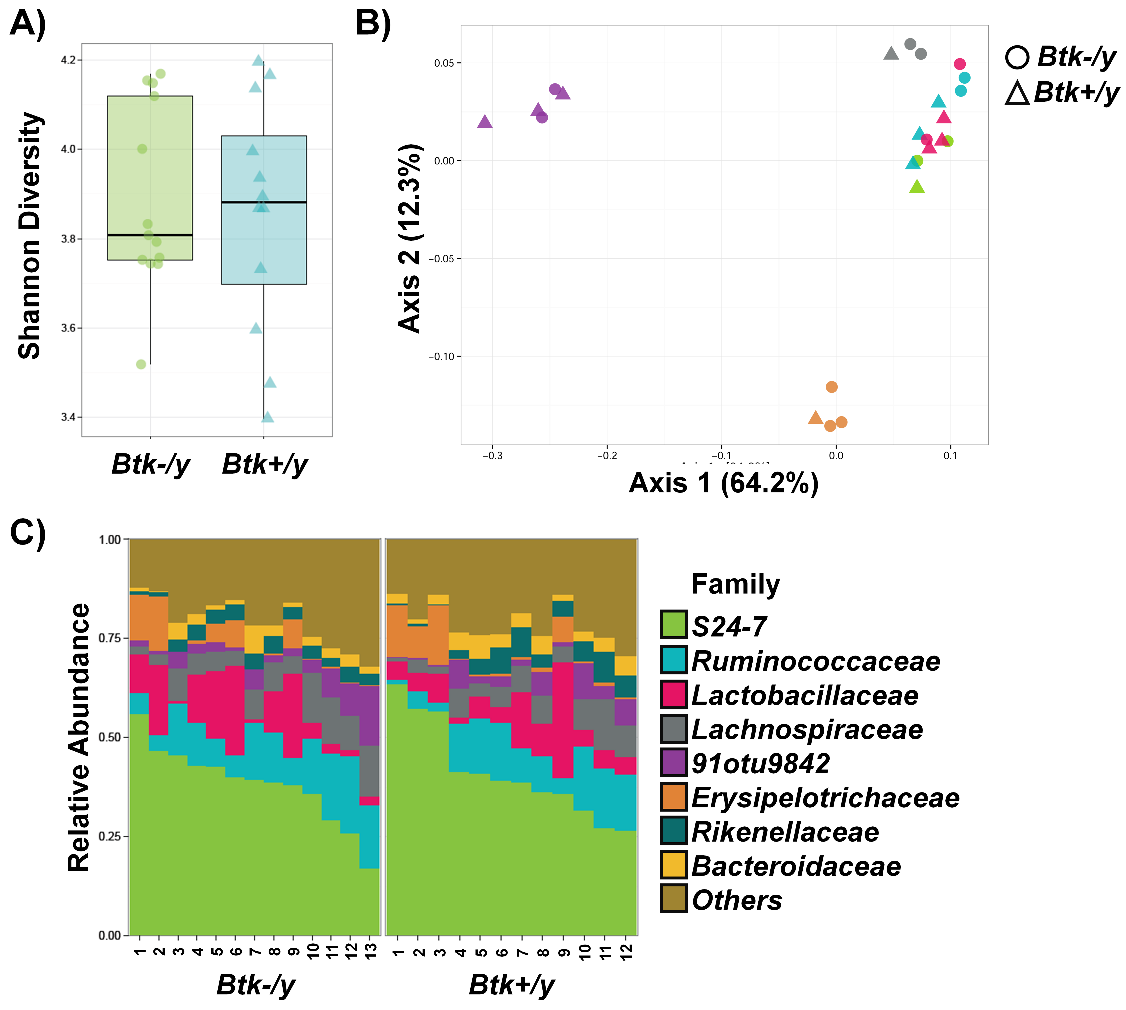

**Supplemental Figure 2. The fecal microbiome is not altered by *Btk*-deficiency in K/BxN mice.** Feces were harvested from 6-8 wk-old *Btk^+/y^* or *Btk^-/y^* K/BxN littermates from n = 6 cages, n = 12-13 mice per group. DNA was isolated from fecal bacteria and microbiome sequencing and analysis was performed. Filtered sequence reads per sample ranged from 208,522 to 312,328. (A) Shannon (alpha) diversity is shown for *Btk^-/y^* (green) and *Btk^+/y^* (blue) K/BxN mice. (B) Individual *Btk^-/y^* (circles) and *Btk^+/y^* (triangles) K/BxN mice are plotted based on principle component analysis. Each color represents a different cage of mice. (C) Family relative abundance is shown for individual mice in each group.
